# A new target of multiple lysine methylation in bacteria

**DOI:** 10.1101/2024.05.15.594293

**Authors:** Shori Inoue, Shogo Yoshimoto, Katsutoshi Hori

## Abstract

The methylation of ε-amino groups in protein lysine residues is known to be an important posttranslational modification in eukaryotes. This modification plays a pivotal role in the regulation of diverse biological processes, including epigenetics, transcriptional control, and cellular signaling. Although less studied in prokaryotes, recent research has begun to reveal the potential role of methylation in modulating bacterial immune evasion and adherence to host cells. In this study, we analyzed the cell surface proteins of the toluene-degrading bacterium *Acinetobacter* sp. Tol 5 by label-free liquid chromatography‒mass spectrometry (LC‒MS) and found that the lysine residues of its trimeric autotransporter adhesin (TAA), AtaA, are methylated. Over 130 lysine residues of AtaA, consisting of 3,630 amino acids and containing 232 lysine residues, were methylated. We identified the outer membrane protein lysine methyltransferase (OM PKMT) of Tol 5, KmtA, which specifically methylates the lysine residues of AtaA. In the KmtA-deficient mutant, most lysine methylations on AtaA were absent, indicating that KmtA is responsible for the methylation of multiple lysine residues throughout AtaA. Bioinformatic analysis revealed that the OM PKMT genes were widely distributed among gram-negative bacteria, including pathogens with TAAs that promote infectivity, such as *Burkholderia mallei* and *Haemophilus influenzae*. Although KmtA has sequence similarities to the OM PKMTs of *Rickettsia* involved in infectivity, KmtA-like PKMTs formed a distinct cluster from those of the *Rickettsia* type according to the clustering analysis, suggesting that they are new types of PKMTs. Furthermore, the deletion of Tol 5 KmtA led to an increase in AtaA on the cell surface and enhanced bacterial adhesion, resulting in slower growth.

**Significance:** Methylation of lysine residues is a posttranslational modification that plays diverse physiological roles in eukaryotes. In prokaryotes however, lysine methylation has been studied only in a limited number of pathogenic bacteria. In this study, we found novel lysine methylation across multiple residues of an outer membrane protein and its methyltransferase, KmtA, in a bacterium from activated sludge. KmtA, along with rickettsial outer membrane protein lysine methyltransferases, which are known to be involved in bacterial pathogenicity, exists in many species of gram-negative bacteria. This finding suggests that methylations are ubiquitous in prokaryotes and are involved in a variety of functions, offering potential strategies for controlling bacterial infections and enhancing the functions of beneficial bacteria for biotechnological applications.

## Introduction

Bacterial fiber proteins on the cell surface are essential mediators of numerous biological functions, including cell adhesion, biofilm formation, motility, DNA transfer, and interaction with the host cell (1–4). These proteins are critical for bacterial survival and cause infection and contamination of pathogenic bacteria (5). On the other hand, fiber proteins from beneficial bacteria have been used for developing various technologies, such as cell-surface display, cell immobilization, bioremediation, and self-repairing materials (6–8).

The intricate structure and composition of these fiber proteins enable a variety of biological functions. Flagella and pili are the major fiber proteins present in a wide range of bacteria. These proteins are composed of thousands of subunits, and their assembly is governed by complex, highly regulated genetic and biochemical processes that ensure their structural integrity and functions (9–11). In addition, many gram-negative bacteria have autotransporter adhesins, which are secreted by the type V secretion system and consist of a passenger domain that is displayed on the outer membrane (OM) and a transmembrane domain that functions as a translocator (12).

The toluene-degrading bacterium *Acinetobacter* sp. Tol 5 exhibits autoagglutination and extremely high adhesiveness to various types of solid surfaces and has been studied for application as an immobilized whole-cell catalyst (13, 14). The high adhesiveness of Tol 5 is mediated by its fibrous cell surface protein AtaA, which is a member of the trimeric autotransporter adhesin (TAA) family, a subtype of autotransporter, and the polypeptide chains of AtaA, which consists of 3,630 amino acids, form a very large homotrimeric fibrous structure 260 nm in length (14). Recent studies have revealed the adhesion preference and force of Tol 5 and the molecular structure and functional domains of AtaA (7, 15–17). On the other hand, electron microscopy has revealed the presence of various fiber proteins on the cell surface of Tol 5 other than AtaA (18). However, a comprehensive analysis of these fiber proteins constituting the cell surface of Tol 5 has yet to be conducted.

In this study, we analyzed the proteome of Tol 5 cells using liquid chromatography‒ mass spectrometry (LC‒MS) and successfully identified a variety of fiber proteins. LC‒ MS analysis coincidentally revealed that the lysine residues of AtaA, the most abundant fiber protein on the cell surface of Tol 5, were methylated. Protein lysine methylation is mainly studied in eukaryotic cells, where it plays diverse biological roles. Although lysine methylation is relatively rare in prokaryotic cells, recent studies have suggested that the methylation of multiple lysine residues on cell surface proteins in prokaryotes, such as *Rickettsia* and *Salmonella*, plays crucial roles in the host immune response during infection and adhesion to host cells (19, 20). Nonetheless, there have been no reports of lysine methylation in TAA family proteins. Therefore, in this study, we focused on the AtaA methylation phenomenon, identified the enzyme responsible for the methylation, and examined the effects of the methylation on the biogenesis of AtaA and physiology of Tol 5.

## Results

### Identification of fiber proteins on the Tol 5 cell surface

We performed label-free LC‒MS analysis on trypsin-digested peptides from Tol 5 cells, and detected 1,977 proteins. These proteins included previously reported cell surface fiber proteins such as AtaA (BCX71826.1) and the Fil pilus subunit FliA (BCX72531.1) (Fig. 1, Table S1) (18). In addition, we detected a protein (BCX73383.1) composed of 3,589 amino acids with sequence similarity to the biofilm-associated protein Bap, which is secreted onto the cell surface by the type 1 secretion system and is involved in biofilm formation (21). The chaperone and usher of F17-like pilus (BCX73228.1, BCX73229.1) were also detected but not its main subunit (BCX73227.1). In *Acinetobacter baumannii,* the Csu pilus was reported to facilitate abiotic surface adherence and biofilm maturation (22), but proteins with sequence similarity to those of the Csu pilus (BCX73208-BCX73213) were not detected in Tol 5.

**Figure 1.**
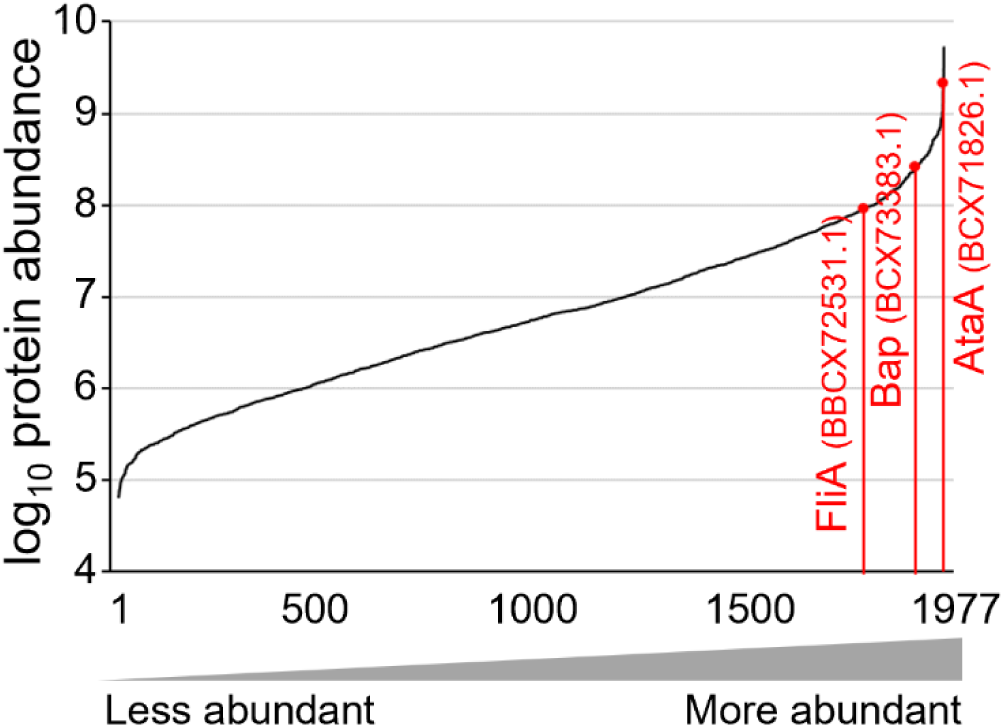
Ranked protein abundance in Tol 5 cells. All quantified proteins were plotted based on quantitative label-free proteomics.

In addition to the identification of cell surface fiber proteins, LC‒MS analysis revealed that the lysine residues of AtaA, the most abundant fiber protein on the cell surface of Tol 5, were methylated (Table S2). Although a few methylated lysine residues were identified in other proteins (Table S2), AtaA was unique in having more than 100 monomethylated lysine residues. The abundances of individual lysine residues and their methylation ratios were determined from the detected peptide fragments (Fig. 2, Table S3, S4). In the analysis, 183 of the 232 lysine residues in AtaA were detected, and methylation was detected at 131 of these residues. The predominant modification was mono-methylation, where a single methyl group is transferred to the ε-amino group of the lysine residues. The monomethylated lysine residues were distributed throughout the AtaA polypeptides. However, dimethylation, where two methyl groups are added, was detected at only 3 residues (Lys616, Lys917, and Lys1218) and at low abundances. Trimethylated lysine residues were not detected.

**Figure 2.**
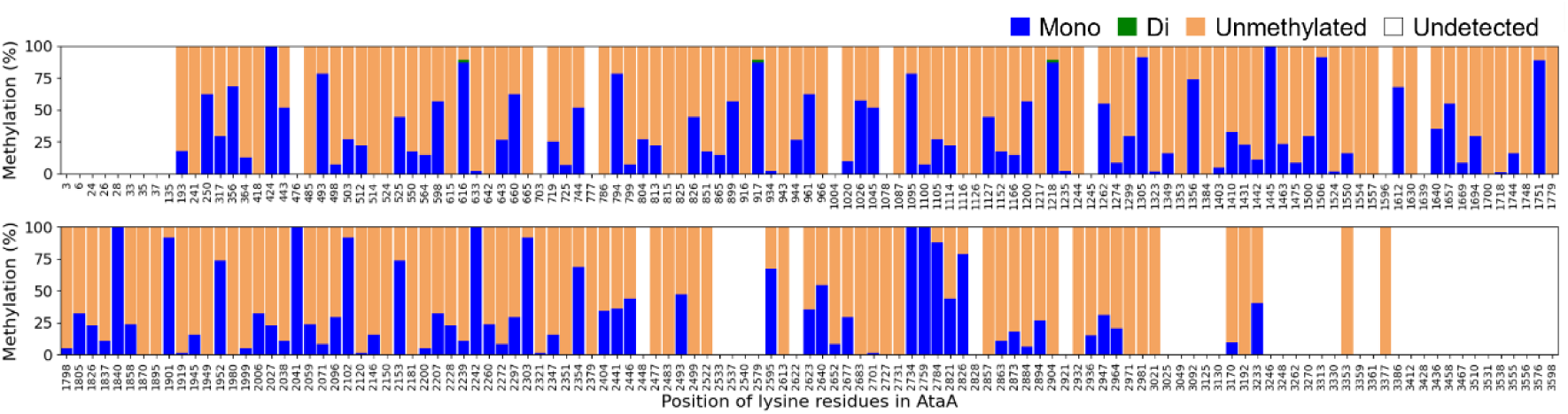
Lysine methylation of AtaA in *Acinetobacter* sp. Tol 5. The percentage of lysine residues that were unmethylated, monomethylated or dimethylated. No trimethylated lysine residues were detected. The data are expressed as the means (n = 2).

### Identification of the enzyme responsible for the methylation of AtaA

To identify the genes involved in the methylation of AtaA, whole-genome databases of Tol 5 (AP024708, AP024709) were analyzed using BLAST. We found a gene encoding an amino acid sequence partially similar to that of OM protein lysine methyltransferases (PKMTs) that were reported to methylate the OM protein OmpB in *Rickettsia* (23, 24). The amino acid sequence of the protein encoded in the identified gene (BCX72587.1) was aligned with those of *Rp*PKMT1 and *Rt*PKMT2 from *Rickettsia* (24) (Fig. 3A). The N-terminal region (Met1-Ile295) of the identified protein, including the methyl group-donor *S*-adenosylmethionine (AdoMet) binding domain (Tyr43-Asn179, Tyr261-Tyr286, and Arg318-Ile331) and the dimerization domain (Thr180-Phe260 and Leu287-Arg317) of *Rp*PKMT1, showed relatively high similarity (34% identity with conserved residues in *Rp*PKMT1 and *Rt*PKMT2), and the residues (Asp102 and His146 in *Rp*PKMT1, Asp74 and His118) of *Rt*PKMT2 that form critical hydrogen bonds with AdoMet were conserved in this new protein, strongly suggesting that it is a PKMT. This new PKMT was named KmtA. On the other hand, the C-terminal region (Asn296-Ala516) of KmtA, including the middle domain (Asn332-Arg447) and the C-terminal domain (Ser448-Val553) of *Rp*PKMT1 showed low similarity (16% identity with conserved residues in *Rp*PKMT1 and *Rt*PKMT2), and Leu103 and Ile130 in the AdoMet binding site of *Rp*PKMT1 (corresponding to Leu75 and Ile102 in *Rt*PKMT2), of which mutation to Ala affects the *K_m_* of OmpB, were not conserved in KmtA.

**Figure 3.**
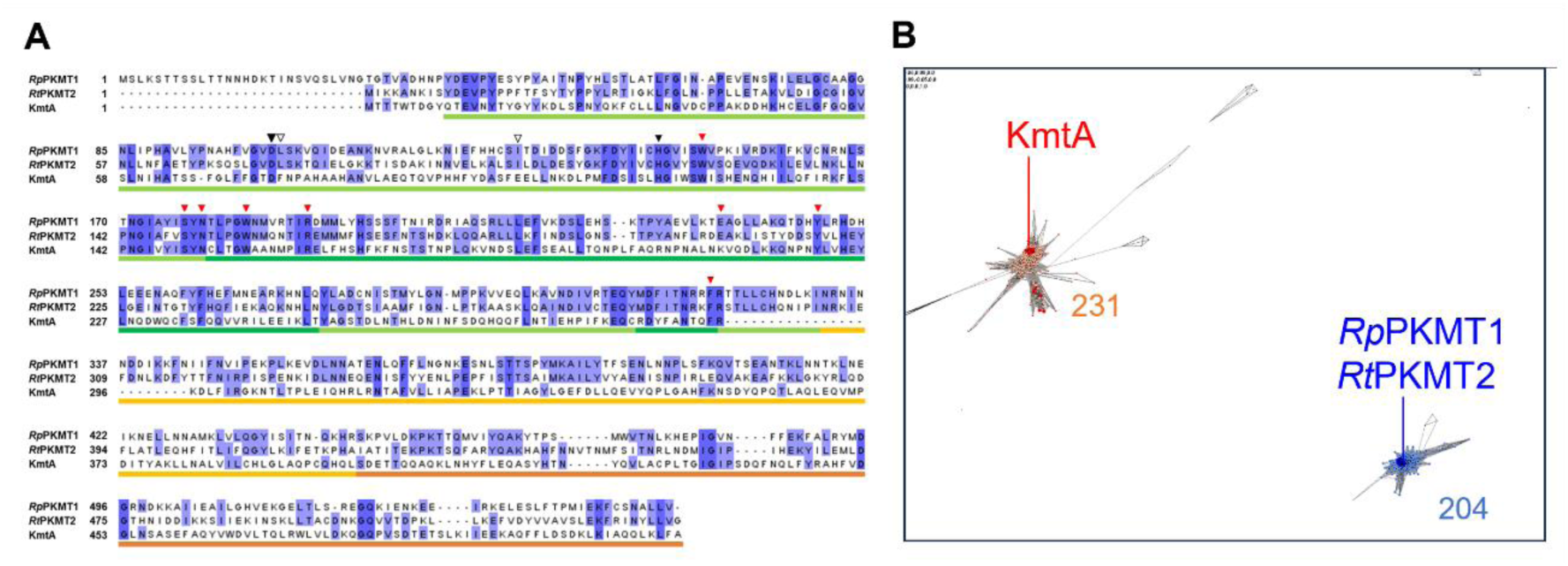
Amino acid sequence alignment and CLANS clustering analysis of *Rp*PKMT1, *Rt*PKMT2 and KmtA. (A) Multiple sequence alignments were obtained using ClustalW (39) and Jalview version 2.11.3.2 (40). Identical residues are highlighted in blue. The bars on the alignments show the structural domains of *Rp*PKMT1 and *Rt*PKMT2 (light green; AdoMet binding domain, green; dimerization domain, yellow; middle domain, orange; C-terminal domain). The black triangles show the residues that form critical hydrogen bonds with AdoMet in *Rp*PKMT1. The white triangles show the residues at which the mutation to Ala affects the Km of OmpB in *Rp*PKMT1. Red triangles show the residues corresponding to the hydrophobic pocket of *Rp*PKMT1 and *Rt*PKMT2 that bind to OmpB. (B) Each dot represents amino acid sequences similar to that of KmtA, that is, OM PKMT. The cluster including KmtA (Cluster A) is indicated by the orange dots. The group including *Rp*PKMT1 and *Rt*PKMT2 (Cluster R) is indicated by light blue dots. Sequences from the reference genomes of *Acinetobacter* and *Rickettsia* are indicated by red and blue dots, respectively. Lines connecting the dots indicate the sequence similarity relationship at the BLAST P value cutoff of 10^-60^.

The amino acid sequences with similarity to that of KmtA were collected using PSI-BLAST and clustered based on sequence similarity using CLANS (25). As shown in Figure 3B, two distinct clusters were observed: Cluster A contained KmtA, and Cluster R contained *Rp*PKMT1 and *Rt*PKMT2. While both clusters contained proteins from a wide range of species across phyla, some phyla were unique to each cluster (Table S5). For example, interestingly, many species in Cyanobacteriota appear only in Cluster A, while many species in PVC (Planctomycetes, Verrucomicrobia and Chlamydiae) group bacteria appear only in Cluster R. For Pseudomonadota (Proteobacteria), while sequences from the same order appear in both clusters, certain sequences are unique to one cluster, such as Rhodobacterales and Cardiobacteriales in Cluster A. Note that the trends observed are based on the limited sequence information currently available. For the genera *Acinetobacter* and *Rickettsia*, homologous sequences were exclusively present in only the respective clusters (Fig. 3B). These results suggest that PKMTs with KmtA-like sequences are distributed in a wide range of bacterial species and are distinct from previously reported OM PKMTs, such as *Rp*PKMT1 and *Rt*PKMT2.

The type strains of *Acinetobacter baumannii*, ATCC 19606 and CCUG 19096, did not possess PKMTs, whereas other pathogenic *Acinetobacter* strains, such as *A. haemolyticus* and *A. junii,* had KmtA-like sequences in their genomes. Furthermore, complete genome sequences with genes encoding OM PKMT were collected and searched for TAA genes, revealing that 27 of the 142 genomic sequences also encode TAA genes (Table S6). Notably, bacteria such as *Burkholderia mallei*, *Burkholderia pseudomallei*, *Haemophilus influenzae*, and *Neisseria meningitidis*, known for their TAAs that mediate bacterial infection, also possess OM PKMT genes (26).

Lysine methylation of AtaA was examined in a *kmtA* gene knockout mutant of Tol 5 (Δ*kmtA*), which does not express KmtA due to point mutations introducing stop codons in the *kmtA* gene. In the LC‒MS analysis, only a single methylated lysine residue was detected in AtaA from Δ*kmtA* (Fig. 4, Table S3, S4). Furthermore, complementation of the *kmtA* gene restored the multiple lysine methylation of AtaA (Fig. 4, Table S3, S4). The average percentage of lysine methylation in AtaA from Δ*kmtA* pKmtA was greater (40%) than that in AtaA from the wild type strain of Tol 5 (WT) (26%). These results indicate that KmtA is responsible for the multiple lysine methylation of AtaA.

**Figure 4.**
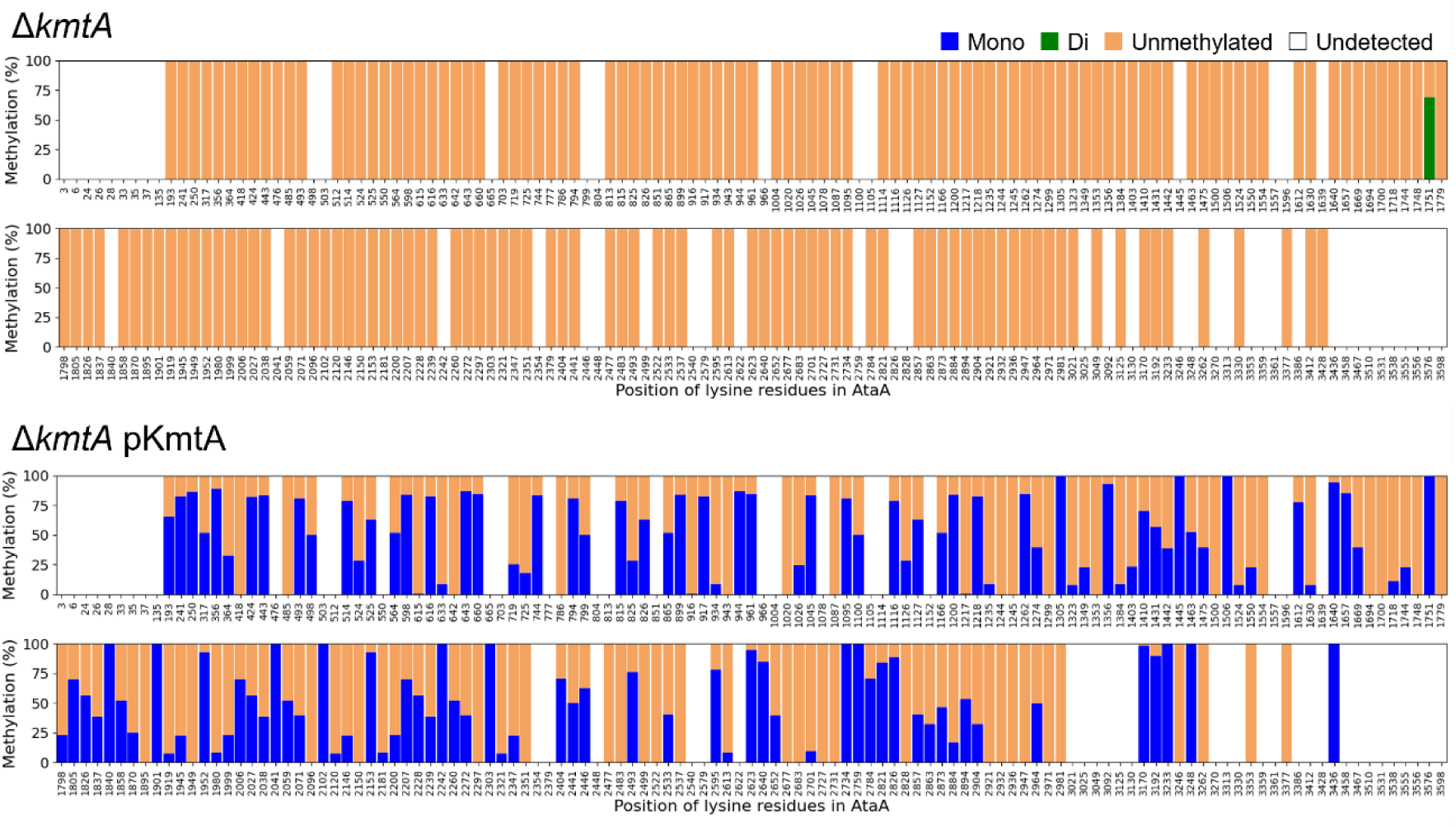
Lysine methylation of AtaA in the mutants of Tol 5. The percentage of lysine residues that were unmethylated, monomethylated or dimethylated. No trimethylated lysine residues were detected. The data are expressed as the means (n = 2).

In the WT and Δ*kmtA* of Tol 5, a small number of proteins other than AtaA with mono-, di-, and trimethylated lysine residues were detected, including the ribosomal protein L11, which is known for multiple lysine methylation (27). In the Δ*kmtA*, the total number of methylated lysine residues in proteins other than AtaA was similar to that in WT (Fig. S1, Table S2), suggesting that the methylation of these proteins is catalyzed by enzymes other than KmtA. In the Δ*kmtA* pKmtA strain, an increase in the number of monomethylated proteins other than AtaA was observed. In this plasmid-complemented strain, the expression level of *kmtA* may have increased due to its increased copy number and enhanced methylation of lysine residues in both AtaA, the original substrate, and other proteins.

### Effects of KmtA on the biogenesis of AtaA and cell physiology

We next investigated whether lysine methylation affects the biogenesis of AtaA fibers in Tol 5. To examine the protein production of AtaA, whole-cell lysates of WT, Δ*ataA*, and Δ*kmtA* were separated by sodium dodecyl sulfate‒polyacrylamide gel electrophoresis (SDS‒PAGE). On the gel stained with CBB, AtaA polypeptides from WT and Δ*kmtA* were detected to have almost the same signal intensity and electrophoretic mobility (Fig. 5A). Considering that the addition of a single methyl group to all 232 lysine residues of AtaA increases its molecular weight by only 3.1 kDa, it is reasonable that methylation of AtaA with a molecular weight greater than 300 kDa would hardly change the electrophoretic mobility of AtaA. To examine the cell surface display of AtaA, WT, Δ*ataA*, and Δ*kmtA* were stained with anti-AtaA antiserum and Alexa 488-conjugated anti-rabbit antibody and then subjected to confocal laser scanning microscopy (CLSM) and flow cytometry. Both WT and Δ*kmtA* exhibited fluorescence, but the intensity of the fluorescence in the Δ*kmtA* was greater than that in WT, indicating that the disruption of the *kmtA* gene enhances the cell surface display of AtaA (Fig. 5B, C).

**Figure 5.**
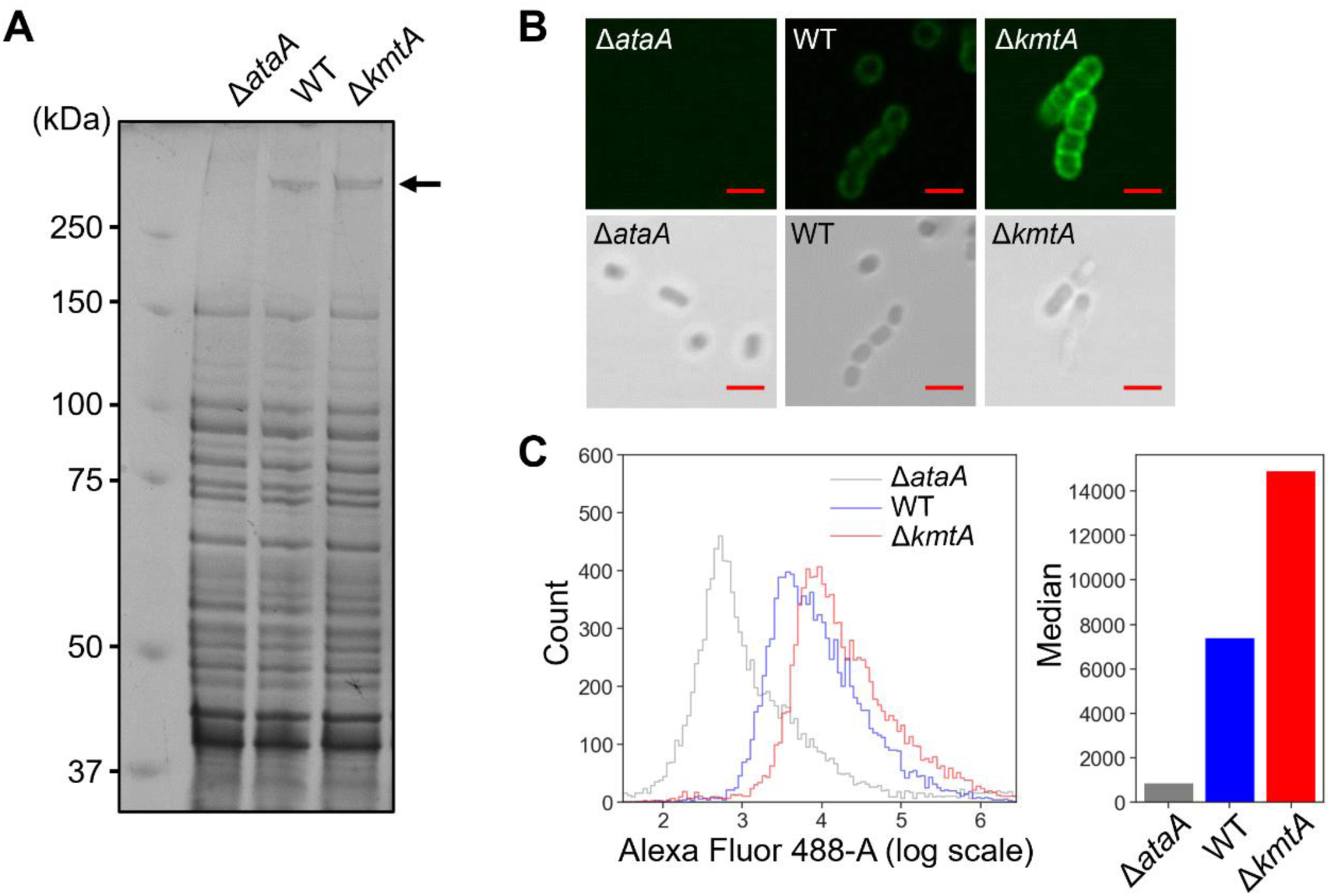
Expression and cell surface display of AtaA in the Tol 5 Δ*kmtA* mutant. (A) CBB staining of Tol 5, its Δ*ataA* mutant, and its Δ*kmtA* mutant cell lysates fractionated according to their molecular weight by SDS‒PAGE. The arrowhead shows the position of the AtaA bands. (B) Immunofluorescence microscopy of Tol 5 and its mutants using an anti-AtaA antiserum and an Alexa-Fluor^®^ 488-conjugated anti-rabbit antibody. Fluorescence and bright fields are shown. Scale bar: 2 µm. (C) Fluorescence flow cytometry analysis of Tol 5 and its mutants using an anti-AtaA antiserum and an Alexa-Fluor^®^ 488-conjugated anti-rabbit antibody.

Subsequently, we examined the adhesiveness and autoagglutination of Tol 5 Δ*kmtA*. Tol 5 Δ*kmtA* showed higher adhesiveness to both hydrophobic polystyrene (PS) plates and hydrophilic glass plates than did WT (Fig. 6A). While both WT and Δ*kmtA* exhibited autoagglutination, the autoagglutination process of Δ*kmtA* was faster than that of WT (Fig. 6B). These results suggest that the KmtA-mediated methylation of AtaA reduces the amount of AtaA on the Tol 5 cell surface, resulting in a decrease in the adhesiveness and autoagglutination of Tol 5 cells.

**Figure 6.**
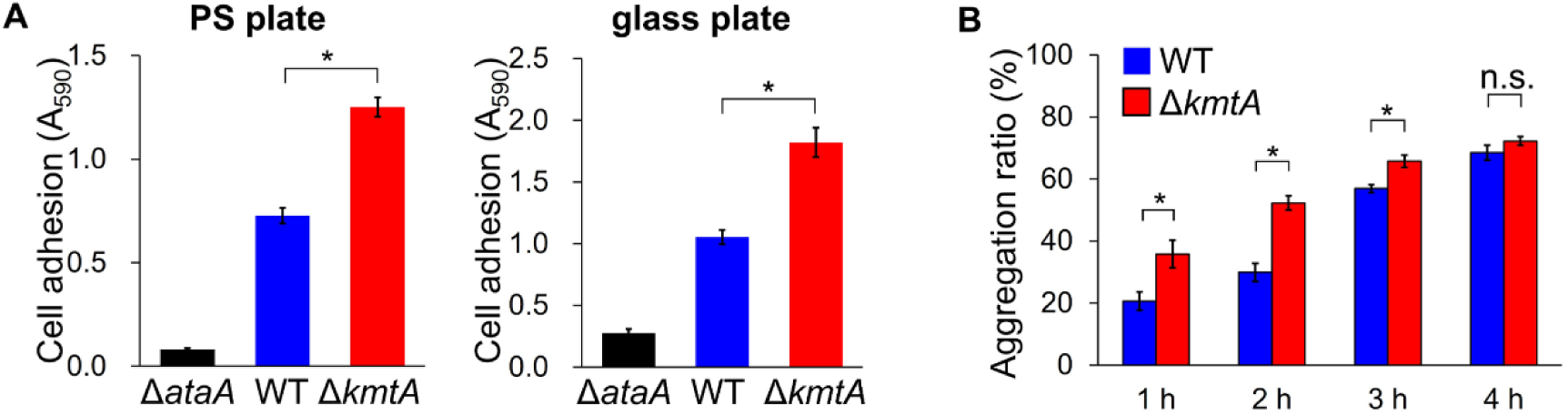
Adherence and autoagglutination assays of the Tol 5 Δ*kmtA* mutant. (A) Adherence assay of Tol 5, its Δ*ataA* mutant and its Δ*kmtA* mutant using 96-well plates made of PS and glass. The data are expressed as the mean ± SEM (n = 5). Statistical significance, \**P* < 0.01. (B) Autoagglutination assay of Tol 5 and its Δ*kmtA* mutant. The graph bars show the autoagglutination ratio (%) at each time point. The data are expressed as the mean ± SEM (n = 3). Statistical significance, \**P* < 0.05; n.s. = not significant.

We further examined the effect of *kmtA* gene disruption on the growth of Tol 5 cells under several conditions. In Luria–Bertani (LB) medium, which inhibits the adhesion and autoagglutination of Tol 5 cells mediated by AtaA (28), WT and Δ*kmtA* exhibited similar growth curves (Fig. 7A). In basal salt (BS) media supplemented with lactate or toluene as the sole carbon source, where Tol 5 cells exhibit high adhesiveness and autoagglutination mediated by AtaA, the growth rate of Δ*kmtA* was lower than that of WT (Fig. 7B, C). In addition, Tol 5 Δ*ataA*Δ*kmtA* showed no significant change in growth rate compared with Tol 5 Δ*ataA* in either LB or BS media (Fig. S2). These results indicate that KmtA promotes the growth of Tol 5 through the methylation of AtaA in environments where cells exhibit adhesiveness and autoagglutination by AtaA.

**Figure 7.**
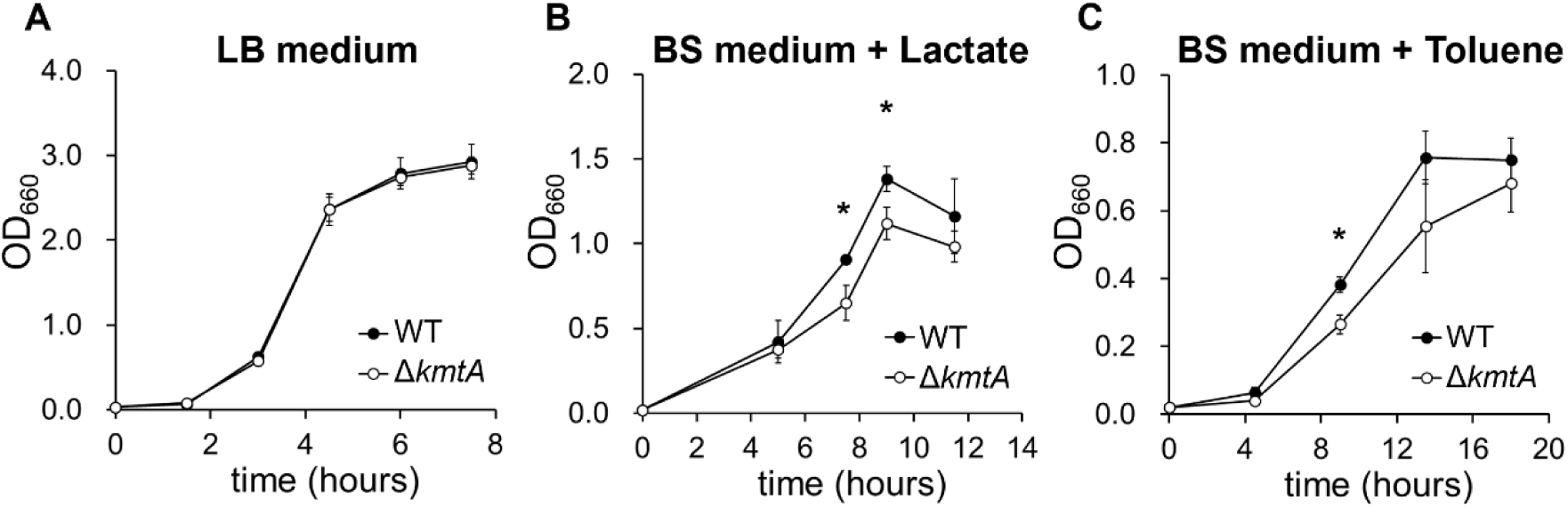
Comparison of the growth of *Acinetobacter* sp. Tol 5 Δ*kmtA* mutant and Tol 5 WT. (A) Tol 5 and its Δ*kmtA* mutant were grown in LB medium. The optical density at 660 nm (OD_660_) of the cell culture was measured. The data are expressed as the mean ± SD (biological replicate n = 3). (B, C) Tol 5 and its Δ*kmtA* mutant were grown in BS media supplemented with lactate or toluene as a carbon source. Polyurethane foam was used as a carrier for cell adhesion. Cells adhering to the polyurethane foam were detached using casamino acid followed by mixing prior to each OD_660_ measurement. The data are expressed as the mean ± SD (biological replicate n = 3). Statistical significance, \**P* < 0.05.

## Discussion

Protein lysine methylation is a common posttranslational modification that occurs in histones and various nonhistone proteins. Lysine methylation plays a crucial role in diverse biological processes, such as the regulation of chromatin conformation, signal transduction, the DNA damage response, protein folding, metabolism, and cell growth in eukaryotes (29, 30). Recent studies have revealed that methylation of cell surface proteins or secreted proteins in pathogenic bacteria contributes to their infectivity (19, 20, 31). However, most studies have focused on histones in eukaryotes, and the biological functions of lysine methylation in bacteria remain largely unknown (32), even though lysine methylation was first discovered in bacteria (33). In this study, we conducted proteome analysis of Tol 5 and coincidentally found the multiple lysine methylation of AtaA (Table S2, S3). This is the first report demonstrating the posttranslational methylation of TAAs and the most extensively methylated case among reported methylations of bacterial OM proteins. Further experiments with Tol 5 Δ*kmtA* revealed that KmtA is responsible for the methylation of AtaA and has a role in controlling the amount of AtaA displayed on the cell surface and the adhesion and agglutination nature of Tol 5 cells, contributing to enhanced growth.

The suppression of the growth of Tol 5 Δ*kmtA* was observed only when a basal salt medium was used for cultivation, while no difference was observed in LB medium (Fig. 7A-C). In a previous study, we reported that the adhesion and autoagglutination of Tol 5 through AtaA were inhibited in LB medium (28). Given that adherent or agglutinating cells often exhibit slower growth than planktonic cells due to the restricted transport and diffusion of cells and substrates (34), the high adhesiveness and autoagglutination of Tol 5 Δ*kmtA* may result in slower growth in BS media, where Tol 5 exhibits high adhesiveness and autoagglutination; this suggests that methylation of AtaA indirectly promotes Tol 5 cell growth by decreasing the amount of AtaA on the cell surface. In fact, we previously reported that Tol 5 displays AtaA fibers poorly in the early growth phase for efficient and fast cell growth (35).

OM proteins other than AtaA in Tol 5 were barely methylated (Table S2), indicating that methylation by KmtA is more specific than that of previously reported rickettsial PKMTs, which methylate not only the main target OmpB but also other OM proteins, such as OmpA and Sca1 (19). Seven of the eight amino acid residues corresponding to the hydrophobic pocket of *Rp*PKMT1 and *Rt*PKMT2 that bind to OmpB were conserved in KmtA (24). On the other hand, L103 and I130 near the AdoMet binding site, which are important for OmpB recognition by *Rp*PKMT1 and *Rt*PKMT2, were not conserved in KmtA. These findings suggest that the substrate recognition mechanism of KmtA is somewhat different from that of rickettsial PKMTs. Clustering analysis further revealed that OM PKMTs with KmtA-like sequences form a distinct cluster separate from rickettsial PKMTs. Comparison of multiple sequence alignments (MSAs) of amino acid sequences derived from each cluster revealed distinct differences between sequences in Clusters A and B (Fig. S3). Notably, the conserved motif of charged amino acids in the N-terminal region of Cluster A proteins (Lys282-Arg295 in KmtA) markedly differs from that in Cluster R proteins. The fact that this region is located within the cleft predicted to bind substrate peptides, according to the working model based on crystal structure analyses of rickettsial PKMTs, may have significant implications. The C-terminal region of OM PKMTs contains several subregions that share sequence similarity within each cluster (Fig. S3). These features suggest that there are two distinct subfamilies of OM PKMTs; thus, we propose to name them OM PKMT A, which includes *Acinetobacter* KmtA, and OM PKMT R, which includes *Rp*PKMT1 and *Rt*PKMT2.

TAAs from pathogenic bacteria have diverse functions, such as adhesion, autoagglutination, biofilm formation, serum resistance, and actin-based motility, and are crucial for infection (36–38). In this study, OM PKMTs were identified in a wide range of bacteria, and 19% of them also possessed TAAs (Table S6). In these bacteria, TAAs may be methylated, similar to AtaA, and this methylation may be involved in diverse functions related to their pathogenicity and biofilm formation. However, some species possess OM PKMTs but not TAAs; OM PKMTs in these species may methylate other OM proteins and contribute to their biological functions, such as growth or infectivity.

In conclusion, we discovered the multiple lysine methylation on AtaA and identified the enzyme responsible for this methylation. This study constitutes the first report of methylation among TAAs. Our findings contribute to a better understanding of posttranslational modifications in bacteria, which is important for both medical and engineering applications.

## Materials and Methods

### Sequence analysis

Amino acid sequence alignments were generated using ClustalW (39) and Jalview version 2.11.3.2 (40). To collect sequences with similarity to KmtA, five iterative PSI-BLAST searches (41) were performed against the uniprot_trembl database (accessed January 20, 2024) at an E-value cutoff of 1e-60. After the highly redundant sequences were removed using HHfilter (41) at a 50% identity cutoff, the sequences with genus-level annotations were extracted. Sequences with similarity to KmtA were also collected from NCBI reference genomes (Assembly level: complete) of *Acinetobacter* species and *Rickettsia* species using BLASTP at an E-value cutoff of 1e-10. These sequences were also analyzed using CLANS (25) at an E-value cutoff of 1e-60. The sequences used in the CLANS analysis were reduced using HHfilter at a 25% identity cutoff and searched against the RefSeq representative genome database of bacteria using tBLASTn (https://blast.ncbi.nlm.nih.gov/Blast.cgi) to collect complete genome sequences containing sequences encoding PKMT. Then, the amino acid sequences of the transmembrane domain of TAAs reported by Thibau A et al., 2019 (26) were searched against the collected complete genome sequences.

### Bacterial strains and growth conditions

The primers, bacterial strains and plasmids used in this study are shown in Tables 1 and 2. *Acinetobacter* sp. Tol 5 and its derivative mutants were grown in basal salt (BS) medium supplemented with toluene or Luria–Bertani (LB) medium at 28 °C as described previously (42). *Escherichia coli* was grown in LB medium at 37 °C. Apramycin (100 µg/ml) and gentamicin (10 µg/ml) were used as needed. Arabinose was added to a final concentration of 0.5% (w/v) for the induction of *kmtA* under the control of the P_BAD_ promoter.

**Table 1.** Strains and plasmids used in this study.

| Strain | Description | Reference |
| --- | --- | --- |
| ***Acinetobacter* sp.** |  |  |
| Tol 5 | Wild type strain | (45) |
| REK123Δ*ataA* | Restriction enzyme-encoding genes and *ataA* knockout mutant of Tol 5 | (44) |
| Δ*kmtA* | *kmtA* gene knockout mutant of Tol 5 | this study |
| Δ*ataA* mutant | Unmarked Δ*ataA* mutant of Tol 5 | (46) |
| Δ*ataA*Δ*kmtA* | *kmtA* gene knockout mutant of Δ*ataA* mutant | this study |
| ***Escherichia coli*** |  |  |
| DH5α | Host for routine cloning | Purchased from Takara (Shiga, Japan) |
| S17-1 | Donor strain for bacterial conjugation | (47) |
| **Plasmid** |  |  |
| pBECAb-apr | For gene knockout | (43) |
| pBECAb-apr-kmtA | For *kmtA* gene knockout | this study |
| pAtaA | Expression vector for AtaA | (14) |
| pKmtA | Expression vector for KmtA | this study |

**Table 2.**
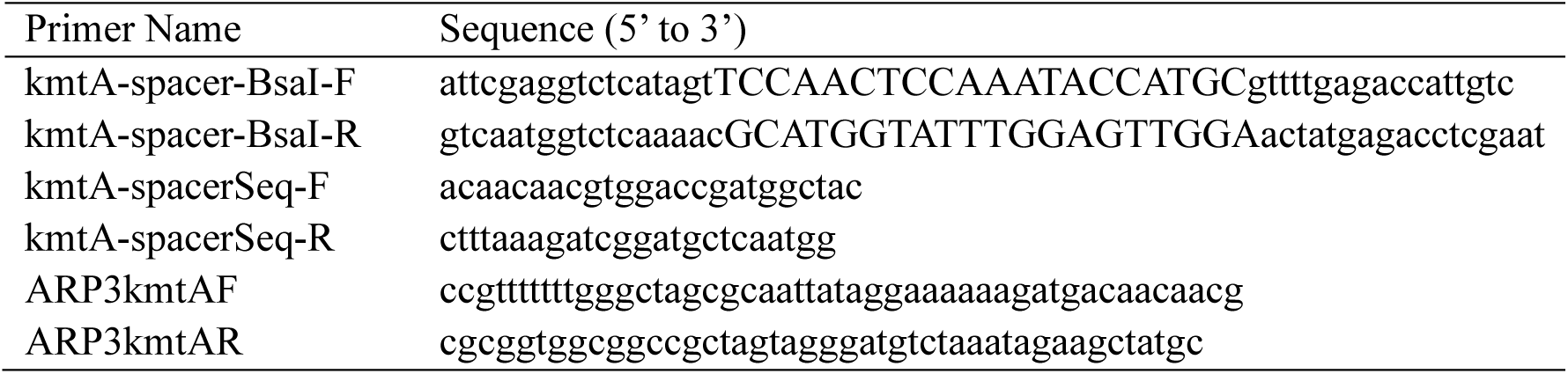
Primers used in this study.

To obtain growth curves, 2 mL of the overnight culture of Tol 5 and its mutants in LB medium was centrifuged (5,000 ×g, 25 °C, 5 min), washed twice with BS medium, and resuspended in BS medium. Twenty microliters of the suspensions were inoculated into 20 mL vials (Maruemu Corporation, Osaka, Japan) containing 2 mL of BS medium or 2 mL of LB medium with 4 pieces of polyurethane foam support with a specific surface area of 37.5 cm^2^/cm^3^ (CFH-30; Inoac Corporation, Nagoya, Japan) in the shape of a rectangle (10 mm × 5 mm × 5 mm). As a carbon source, 8 µL of 50% L-lactate or 2.5 µL of toluene was added to the medium, and the vials were incubated at 28 °C with shaking at 115 rpm. At each time point during cultivation, three vials were collected, and the cultures were mixed vigorously with 100 µL of 10% Casamino Acids technical grade (CA-T; Becton, Dickinson and Company, Franklin Lakes, NJ, USA) to detach the bacterial cells from the support (28). The OD_660_ of the cell suspension was measured by a UV‒Vis spectrophotometer (UV1800; Shimadzu Corporation, Kyoto, Japan).

### Plasmid construction and gene knockout

The *kmtA* gene knockout mutants (Δ*kmtA*) were generated using the cytidine base editing system for *Acinetobacter* (43). The inserted DNA dimer, encoding the sequence of the single guide RNA, was prepared by mixing oligo DNAs (kmtA-spacer-BsaI-F and kmtA-spacer-BsaI-R) in 50 mM NaCl solution and gradually cooling from 95 °C to 18 °C at 0.1 °C/sec. This dimer was introduced into pBECAb-apr using the NEBridge Golden Gate Assembly Kit (New England Biolabs, Ipswich, MA, USA) to generate pBECAb-apr-kmtA. Transformation of Tol 5 and its mutant was performed through REK123Δ*ataA*, following the method reported in Ishikawa M et al. 2023 (44). After overnight culture of Tol 5 and its mutant harboring pBECAb-apr-kmtA in BS medium to promote mutation, the cells were spread on a BS agar plate containing 5% (w/v) sucrose. Mutation of the *kmtA* gene in the resultant colonies was confirmed by sequencing the DNA from colonies via PCR using KOD FX Neo (TOYOBO, Osaka, Japan).

To construct pKmtA, a fragment containing the deduced Shine‒Dalgarno (SD) sequence and the entire *kmtA* gene was amplified by colony PCR from Tol 5 using the primers ARP3kmtAF and ARP3kmtAR. The PCR product was inserted into the EcoRI-XbaI site of pARP3, generating pKmtA. Transformation of Δ*kmtA* was carried out by bacterial conjugation with the donor strain *E. coli* S17-1 harboring pKmtA as previously described (14).

### Sample preparation for mass spectrometry

Sample preparation for LC‒MS analysis was performed as described previously by Engström P et al. 2021 (19). One microliter of the suspension of Tol 5 and its mutants cultured in LB medium was transferred to a 1.5 mL microtube and centrifuged (5,000 ×g, 4 °C, 5 min) to collect the precipitated bacterial cells. The cells were resuspended in 333 µL of Tris-EDTA solution (10 mM Tris-HCl, 10 mM EDTA, pH 7.6), incubated at 45 °C for 90 min and then boiled at 95 °C for 10 min. After cooling to room temperature, 20 μL of 50 mM NH_4_HCO_3_ (pH 7.5) and 50 μL of 0.2% (w/v) RapiGest (Waters Corporation, Milford, MA, USA) diluted in 50 mM NH_4_HCO_3_ were added to 50 µL of the samples and heated at 80 °C for 15 min. After cooling to room temperature, the samples were digested with 1 μg of Pierce Trypsin Protease (Thermo Fisher Scientific, Waltham, MA, USA) at 37 °C overnight. To hydrolyze the RapiGest, 20 μL of 5% (v/v) trifluoroacetic acid was added to the samples, which were then incubated at 37 °C for 90 min. After centrifugation (15,000 ×g, 4 °C, 25 min), the samples were desalted using ZipTip (ZTC18M096; Merck Millipore, Burlington, MA, USA) and dried using a CVE-3100 (Tokyo Rikakikai Co., Ltd., Tokyo, Japan).

### Liquid chromatography–mass spectrometry

The digested peptides were analyzed by nanoflow reverse-phase LC followed by tandem MS using a Q Exactive hybrid mass spectrometer (Thermo Fisher Scientific). The capillary reverse-phase HPLC-MS/MS system was composed of a Dionex U3000 gradient pump equipped with a VICI CHEMINERT valve. Q Exactive was equipped with a nanoelectrospray ionization (NSI) source (AMR). The desalted peptides were loaded into a separation capillary C18 reverse-phase column (NTCC-360/100-3-125, 125 × 0.1 mm; Nikkyo Technos Co., Ltd., Tokyo, Japan). The peptide spectra were recorded over the mass range of m/z 350–1,800 using the Xcalibur 4.1.50 system (Thermo Fisher Scientific). MS spectra were recorded repeatedly, followed by 20 data-dependent high-energy collisional dissociation (HCD) MS/MS spectra generated from the 10 highest intensity precursor ions. MS/MS spectra were interpreted, and peak lists were generated using Proteome Discoverer 2.4.1.15 (Thermo Fisher Scientific). Searches against the *Acinetobacter* sp. Tol 5 whole genome database (AP024708, AP024709; NCBI) were performed using SEQUEST (Thermo Fisher Scientific). The search parameters were set as follows: enzyme selected with two maximum missing cleavage sites, a mass tolerance of 10 ppm for peptide tolerance, 0.02 Da for MS/MS tolerance, and variable modification of oxidation (M), monomethylation (R, K), di-methylation (R, K), trimethylation (R, K), and N-terminal acetylation. Peptide identification was based on significant Xcorr values (high confidence filter). The abundance of individual lysine residues in AtaA was calculated as the sum of the abundance of the peptides containing these residues, excluding peptides with detected modifications of ambiguous localization. For the peptides whose amino acid sequences appeared repeatedly at multiple positions in AtaA, the abundance value was divided by the number of repeats and assigned equally to all of those positions. The percentage of lysine methylation (mono-, di-, tri-, or unmethylated) was calculated by dividing the abundance of each methylated residue by the total abundance of the residue and multiplying by 100.

### Polyacrylamide gel electrophoresis

One milliliter of the suspension of Tol 5 or its mutants was transferred to a 1.5 mL microtube and centrifuged (5,000 ×g, 4 °C, 5 min). The precipitated cells were resuspended in SDS‒PAGE sample buffer (5% (v/v) 2-mercaptoethanol, 2% (w/v) SDS, 0.02% (w/v) bromophenol blue, and 62.5 mM Tris-HCl, pH 6.8) and boiled at 100 °C for 10 min. The samples were analyzed by SDS‒PAGE using a 7.5% polyacrylamide gel, followed by staining with Coomassie Brilliant Blue (CBB).

### Immunofluorescence microscopy

Immunofluorescence microscopy was performed as described previously (42) with slight modifications. Twenty microliters of bacterial cell culture was placed on a glass slide (TF1205M; Matsunami Glass Ind., Ltd., Osaka, Japan) and incubated at room temperature for 15 min. The cells were fixed with an equal volume of 4% paraformaldehyde, incubated at room temperature for 15 min, and washed twice with phosphate-buffered saline containing Tween 20 (PBS-T; 1.7 mM K_2_SO_4_, 5 mM NaHPO_4_, 150 mM NaCl, 0.05% (w/v) Tween 20, pH 7.4). The samples were incubated with anti-AtaA_699-1014_ antiserum (42) at a 1:10,000 dilution in PBS-T at room temperature for 30 min. The samples were washed twice with PBS-T and incubated with an AF488-conjugated anti-rabbit antibody (Cell Signaling Technology, Inc., Danvers, MA, USA) at a 1:1,000 dilution in PBS-T at room temperature for 30 min. The samples were washed three times with PBS-T once with pure water and observed under a confocal laser scanning fluorescence microscope (FV1000; Olympus, Tokyo, Japan).

### Flow cytometry

Flow cytometry was performed as described previously (14) with slight modifications. One milliliter of bacterial culture was transferred to a 1.5 mL microtube and centrifuged (5,000 ×g, 4 °C, 5 min). The precipitated cells were then fixed with 4% paraformaldehyde at room temperature for 15 min. After washing twice with PBS-T containing 1% CA-T, the samples were incubated with anti-AtaA_699-1014_ antiserum at a 1:10,000 dilution in PBS-T containing 1% CA-T at 4 °C for 1 h with inversion mixing. The samples were washed twice with PBS-T containing 1% CA-T and incubated with an AF488-conjugated anti-rabbit antibody at a 1:1,000 dilution in PBS-T containing 1% CA-T at room temperature for 1 h with inversion mixing. The immunostained samples were washed in PBS-T containing 1% CA-T and resuspended in 500 µL of PBS-T, and fluorescence was measured using a flow cytometer (CytoFLEX S; Beckman Coulter, Brea, CA, USA).

### Bacterial adhesion and autoagglutination assay

Adherence assays and autoagglutination assays were performed as previously described (35) with slight modifications. For the adherence assay, the suspensions of Tol 5 and its mutants were placed into a 96-well polystyrene (PS) microplate (353072; Becton, Dickinson and Company) and a 96-well glass plate (FB-96; Nippon Sheet Glass Co., Ltd., Tokyo, Japan), followed by incubation at 28 °C for 2 h. The adhered cells were stained with 0.1% crystal violet, and the stain was eluted from the cells with 200 μl of 70% ethanol. The absorbance at 590 nm (A_590_) of the eluted solution was measured by a microplate reader (ARVO V3; PerkinElmer, Waltham, MA, USA). In the autoagglutination assay, test tubes (TEST13-100NP, Iwaki, Tokyo, Japan) containing 8 ml of cell suspension were left at room temperature for sedimentation of the cell clumps generated by autoagglutination. The autoagglutination ratio was calculated based on the decrease in the OD_660_ value of the cell suspension due to cell sedimentation using the following equation:

Autoagglutination ratio (%) = (initial OD_660_ − OD_660_ after standing)/(initial OD_660_) × 100

## Author Contributions

KH organized the entire study. All the authors designed the study. SY conducted preliminary experiments. SI and SY conducted the bioinformatic analysis. SI conducted all other experiments. All the authors wrote and reviewed the paper.

## Acknowledgments

The authors wish to acknowledge the Molecular Structure Center, Institute of Transformative Bio-Molecules (WPI-ITbM), Nagoya University, for technical support with the LC‒MS analysis. The authors also thank Hiroya Oka for kindly discussing the proteome analysis; Masahito Ishikawa for kindly discussing the gene knockout; Michio Homma for kindly reviewing the manuscript; and Tomoya Karakama, Jin Mishima, and Satoshi Ishii for their technical assistance. This research was supported by the Graduate Program of Transformative Chem-Bio Research at Nagoya University supported by MEXT (WISE Program) to SI and the Japan Society for the Promotion of Science (JSPS) KAKENHI (Grant Numbers JP20H00319, JP21H05227, and JP24H00043) to KH.

