## Supporting Information for "A new target of multiple lysine methylation in bacteria"


**Supporting Information includes:**

Figures S1 to S3
Tables S1 to S6 (Excel files)

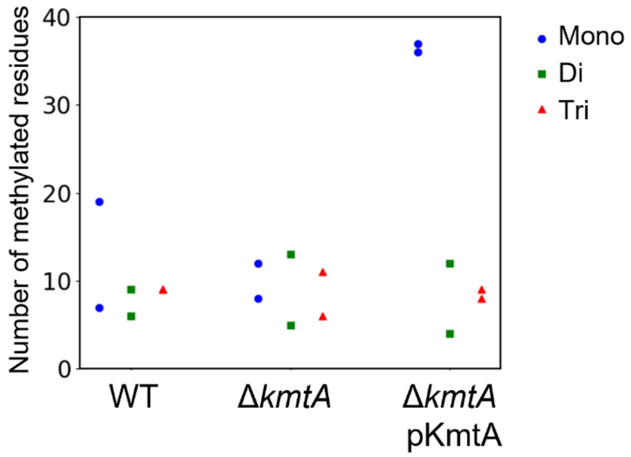


**Figure S1. The number of methylated lysine residues except for residues in AtaA.** Count of methylated lysine residues (mono-, di-, and tri-) in peptides quantified by LC–MS, in which the peptides from AtaA were excluded (biological replicates, n = 2).

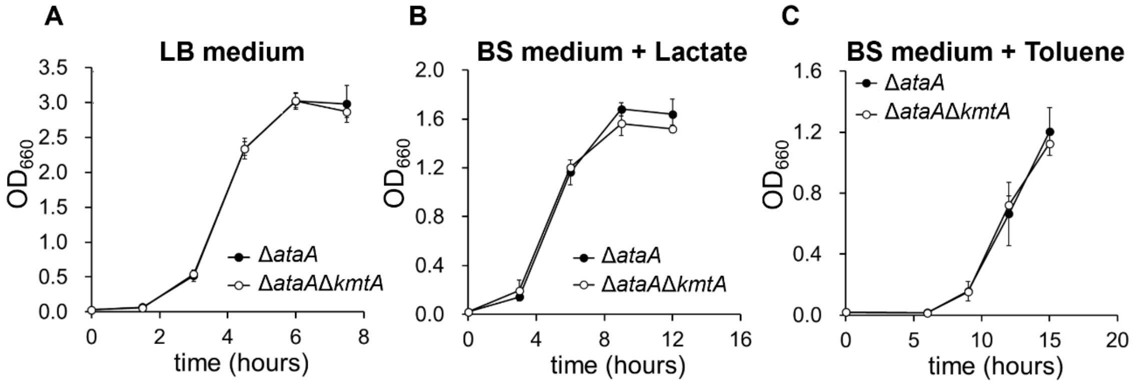


**Figure S2. Growth of *Acinetobacter* sp. Tol 5 Δ*ataA* mutant and Δ*ataA*Δ*kmtA* double knockout mutant**. (A) Δ*ataA* and its Δ*kmtA* mutant were grown in LB medium. The optical density at 660 nm ($OD_{660}$) of the cell culture was measured. The data are expressed as the mean ± SD (biological replicate n = 3). (B, C) Tol 5 Δ*ataA* and its Δ*kmtA* mutant were grown in BS media supplemented with lactate or toluene. Polyurethane foam was used as a carrier for cell adhesion. Cells adhering to the polyurethane foam were detached using casamino acid followed by mixing prior to each $OD_{660}$ measurement. The data are expressed as the mean ± SD (biological replicate n = 3).

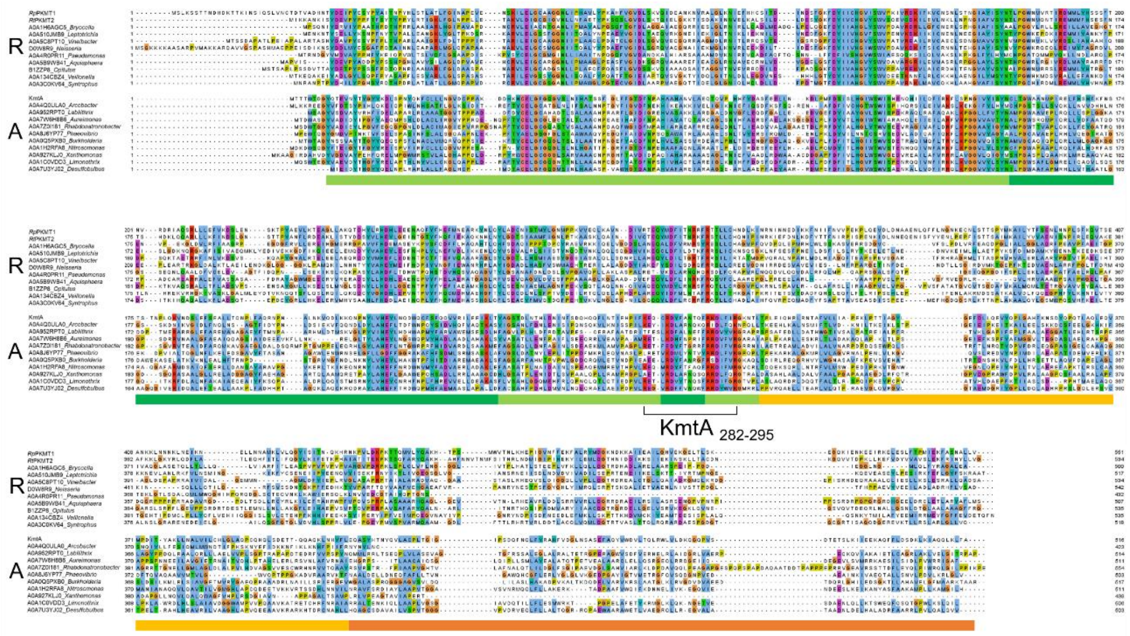


**Figure S3. A comparison of multiple sequence alignments of amino acid sequences derived from Cluster A and Cluster R shown in Figure 3B.** Multiple sequence alignments (MSAs) were obtained using ClustalW (39) and visualized using the clustal coloring method of Jalview version 2.11.3.2 (40). In addition to *Rp*PKMR1, *Rt*PKMT2 and KmtA, 10 sequences were extracted from each cluster (Cluster A and Cluster R) to create the MSAs.

**Table S1. Label-free LC–MS analysis of Tol 5.**

**Table S2. Peptides with methylation.**

**Table S3. Peptides of AtaA.**

**Table S4. Lysine residues in AtaA.**

**Table S5. Genera found in the two clusters in Figure 3B.**

**Table S6. Strains with complete genome sequences including PKMT genes.**
